# Evaluating stably expressed genes in single cells

**DOI:** 10.1101/229815

**Authors:** Yingxin Lin, Shila Ghazanfar, Dario Strbenac, Andy Wang, Ellis Patrick, Dave Lin, Terence Speed, Jean YH Yang, Pengyi Yang

## Abstract

**Background:** Single-cell RNA-seq (scRNA-seq) profiling has revealed remarkable variation in transcription, suggesting that expression of many genes at the single-cell level are intrinsically stochastic and noisy. Yet, on cell population level, a subset of genes traditionally referred to as housekeeping genes (HKGs) are found to be stably expressed in different cell and tissue types. It is therefore critical to question whether stably expressed genes (SEGs) can be identified on the single-cell level, and if so, how their expression stability can be assessed? We have developed a computational framework for ranking expression stability of genes in single cells. Here we evaluate the proposed framework and characterize SEGs derived from two scRNA-seq datasets that profile early human and mouse development.

**Results:** Here, we show that gene expression stability indices derived from the early human and mouse development scRNA-seq datasets are highly reproducible and conserved across species. We demonstrate that SEGs identified from single cells based on their stability indices are considerably more stable than HKGs defined previously from cell populations across 10 diverse biological systems. Our analyses indicate that SEGs are inherently more stable at the single-cell level and their characteristics reminiscent of HKGs, suggesting their potential role in sustaining essential functions in individual cells.

**Conclusions:** SEGs identified in this study have immediate utility both for understanding variation/stability of single-cell transcriptomes and for practical applications including scRNA-seq data normalization, the proposed framework can be applied to identify genes with stable expression in other scRNA-seq datasets.

## Background

A hallmark of single-cell RNA-seq (scRNA-seq) data has been the remarkable variation in gene transcription that occurs at the level of individual cells [1]. The high degree of variation has led to the appreciation that transcription of genes at the single-cell level are comparatively noisier than on the cell population level [2]. Indeed, a subset of genes are thought to be characterized by their stochastic expression [3]. Supporting this notion, genes were found to show transcriptional bursting where their expression varies drastically in individual cells [4, 5]. Furthermore, a large number of genes from scRNA-seq data exhibit bimodality or multimodality of non-zero expression values [6], suggesting that many of these genes may be expressed at different levels in the same and/or different cells. These phenomena illustrate that expression stochasticity is an intrinsic property of many genes on the single-cell level [7].

On the cell population level, however, a subset of genes traditionally referred to as housekeeping genes (HKGs) [8, 9] are found to be stably expressed in different cell types, tissue types and developmental stages [10]. The concept of HKGs is often related to the gene set required to maintain basic cellular functions and therefore is crucial to the understanding of the core transcriptome that is required to sustain life [11,12,13]. Early studies such as those by [14], [15], [8], and [16] were conducted to define HKGs using serial analysis of gene expression (SAGE) or microarrays. With the advent of biotechnologies, follow-up studies using more comprehensive data sources such as those by [17] and [18], and high-throughput RNA sequencing (RNA-seq) by [10] and [19], have refined the list of HKGs from populations of cells.

Taken together, the findings from bulk transcriptome data of cell populations and the stochasticity in gene expression observed in individual cells from scRNA-seq data, several fundamental questions arise including (i) Can patterns of stably expressed genes be identified from single cell data? And if so, (ii) how stable are they across individual cells from different tissue types and biological systems? (iii) What properties do such genes have? And (iv) how do they compare to HKGs defined from bulk transcriptome data?

Leveraging the advances of scRNA-seq techniques [20, 21], we have developed a computational framework to rank genes based on various properties extracted from scRNA-seq data to characterize their expression stability in individual cells [22]. These genes were subsequently utilized for scRNA-seq data normalization and integration. To address the questions posed above, here, we evaluated the reproducibility of the proposed framework on two large-scale high-resolution scRNA-seq datasets in which a wide range of cell types and developmental stages were profiled in human [23] and mouse [24]. We referred to the list of stably expressed genes derived from these two datasets as “hSEG” and “mSEG” for human and mouse respectively, and collectively as “SEGs”. We subsequently evaluated the stability of SEGs on the two scRNA-seq datasets from which they were identified and eight independent scRNA-seq datasets generated from diverse tissues and biological systems, and different sequencing protocols. Compared to HKGs previously defined using bulk microarray [16] or RNA-seq datasets [10], SEGs identified on the single-cell level are considerably more stable in all 10 biological systems, demonstrating the higher resolution enabled by scRNA-seq data for identifying genes that are truly stably expressed across individual cells, and suggesting their potential roles in maintaining essential functions in individual cells.

Our analyses highlight the previously unappreciated gene stability at the single-cell level. Our computational framework also allows further identification and refining of SEGs in other scRNA-seq datasets. This may have broad application in normalization [25, 26] and removal of unwanted variation [27, 28, 22] in scRNA-seq as well as bulk sequencing datasets generated from various experiments.

## Data Description

### scRNA-seq data processing

A collection of 10 publicly available scRNA-seq datasets (Table 1) were utilized in this study. These datasets were downloaded from either NCBI GEO repository or the EMBL-EBI ArrayExpress repository. Fragments per kilobase of transcript per million (FPKM) values or counts per million (CPM) from their respective original publications were used to quantify full length gene expression for datasets generated by SMARTer or SMART-Seq2 protocols. UMI-filtered counts were used to quantify gene expression for the InDrop dataset. All datasets have undergone cell-type identification using biological knowledge assisted by various clustering algorithms from their respective original publications which we retain for evaluation purposes. For each dataset, genes with more than 80% missing values (zeros) were removed, with the remaining genes considered as expressed in that dataset. These filtered datasets were used for all subsequent analyses.

**Table 1.** Summary of scRNA-seq datasets utilized for stably expressed gene identification and/or evaluation in this study.

| ID | Publication | Description | Organism | # cell | # class | Protocol | Purpose |
| --- | --- | --- | --- | --- | --- | --- | --- |
| E-MTAB-3929 | [23] | Early human development | Human | 1529 | 5 | SMART-Seq2 | identify & evaluate |
| GSE45719 | [24] | Early mouse development | Mouse | 269 | 8 | SMART-Seq2 | identify & evaluate |
| GSE94820 | [29] | Peripheral blood mononuclear cells | Human | 1140 | 5 | SMART-Seq2 | evaluate |
| GSE75748 | [30] | PSCs and endoderm progenitors | Human | 1018 | 7 | SMARTer | evaluate |
| GSE72056 | [31] | Multicellular metastatic melanoma | Human | 4645 | 7 | SMART-Seq2 | evaluate |
| GSE67835 | [32] | Adult and fetal brain | Human | 466 | 8 | SMARTer | evaluate |
| GSE60361 | [33] | Cortex and hippocampus | Mouse | 3005 | 7 | SMARTer | evaluate |
| GSE52583 | [34] | Developmental lung epithelial cells | Mouse | 198 | 4 | SMARTer | evaluate |
| E-MTAB-4079 | [35] | Mesoderm diversification | Mouse | 1205 | 4 | SMART-Seq2 | evaluate |
| GSE84133 | [36] | Pancreas inter- and intra-cells | Mouse | 822 | 13 | InDrop | evaluate |

## Analyses

### A computational framework for measuring gene expression stability in single cells

Briefly, the framework (Figure 1A) extracts a set of stability features including λ, σ^2^, α>^*^, and the *F*-statistics (introduced below), and derives a stability index for each gene on the singlecell level.

**Figure 1.**
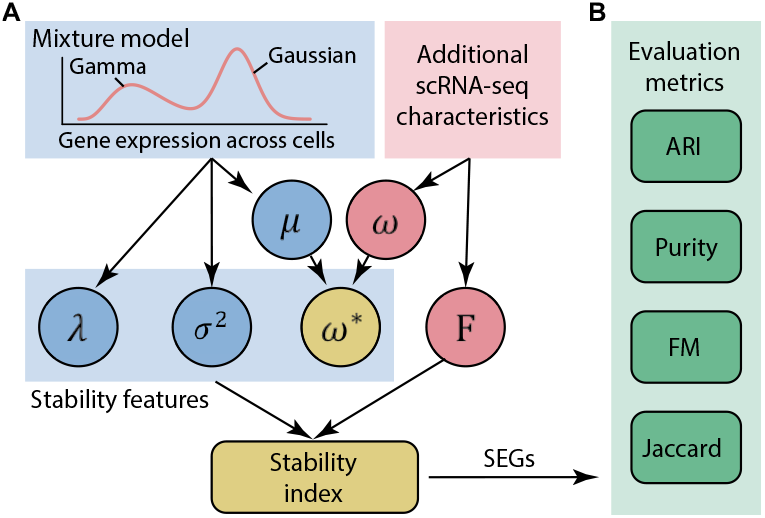
Schematic illustration of the computational framework for deriving gene stability index on the single-cell level. (A) Stability features extracted directly from the mixture model are colored in blue. Those extracted from additional scRNA-seq data characteristics are in red. The overall stability index is derived from the combination of all stability features. (B) Evaluation metrics used for evaluating gene expression stability in scRNA-seq datasets.

The μ and σ^2^ denote the mean and variance of the Gaussian component from fitting a Gamma-Gaussian mixture model [37] to the non-zero expression values of a gene *x* across individual cells. The joint density function *f*(.) of the mixture model is defined as follows:

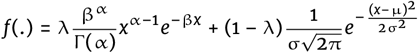

where 0 ≤ λ ≤ 1 is the mixing proportion indicating the proportion of cells in the Gamma component in the fitted model. Genes whose expression profiles are with low mixing proportion (λ) and small variance (σ^2^) are unimodal and relatively invariant across cells and therefore more likely to be stably expressed.

The ω denotes the percentage of zeros of a gene across cells. The measured expression level for a given gene and cell may be zero due to technical dropout, stochastic expression, or no transcription occurring at all for that gene [38]. Thus, SEG would have relatively small ω (i.e. low proportion of zeros), since they are expected to be expressed in all cells. However, lowly expressed genes tend to have a higher proportion of zeros than highly expressed genes simply due to technical dropouts [39]. We therefore regularize ω by average expression level μ in Gaussian component of each gene as ω* = ω · minmax(μ) p>such that we anticipate more dropout events for SEGs with low expression compared to highly expressed genes. When predefined cell type annotation is available for a given dataset, the *F*-statistics can be utilized as another stability feature to select for genes in which we observe the same average gene expression across different pre-defined cell types. Together, genes with small λ, σ^2^, ω* and *F*-statistic are unimodal, expressed with low variance, with relatively low percentage of zeros, and expressed similarly across all cell types, respectively, and are more likely to be stably expressed.

The expression stability index is defined for each gene by combining these four stability features. Specifically, genes are ranked first in increasing order with respect to λ, σ^2^, ω* and *F*-statistics; and the ranks from each stability features are rescaled to range from 0 to 1. The stability index for each gene is defined as the average of its scaled rankings across all four stability features. Thus, genes are ranked in terms of their degree of evidence towards expression stability in individual cells and can be selected by adjusting the stability index threshold. The subsequent evaluation can be conducted to assess the stability and generalization property of selected SEGs in other biological systems using various evaluation metrics (Figure 1B; Section 2.3).

### Genes are reproducibly ranked by their expression stability in single cells

To investigate if some genes are inherently more stable in expression on the single-cell level, we utilized two large-scale high-resolution scRNA-seq datasets to quantify genes that are expressed at steady level across different cell types and developmental stages in early human and mouse development, respectively. Briefly, the two scRNA-seq datasets contain (i) transcriptome profiles of 1,529 individual cells derived from 88 human preimplantation embryos ranging from 3rd to 7th embryonic day [23] and (ii) transcriptome profiles of 269 individual cells derived from oocyte to blastocyst stages of mouse preimplantation development [24] (Table 1). The wide range of cell types and developmental stages [40] captured by early mammalian development such as in these two datasets provide a suitable starting point for identifying SEGs that may allow generalization to various cell/tissue types and biological systems.

We first looked at the proportion of zeros per gene across all profled cells in the early human and mouse development scRNA-seq datasets respectively. We found that a large percentage of genes have more than 50% zero quantification across cells in both datasets (Figure 2A), suggesting most of the genes are transiently expressed during different developmental stages in both human and mouse. We observed that the mixing proportion of each gene, the variance and mean expression level from the Gaussian component from the mixture model, and the F-statistics calculated using pre-defined cell labels were different in the two scRNA-seq datasets (Figure 2B). This suggests the need to calculate gene expression stability in early human and mouse development separately. Nevertheless, by combining the scaled ranks of genes with respect to each stability feature, the stability index distributions derived for human and mouse genes appeared to be highly comparable (Figure 2B, bottom right panel).

**Figure 2.**
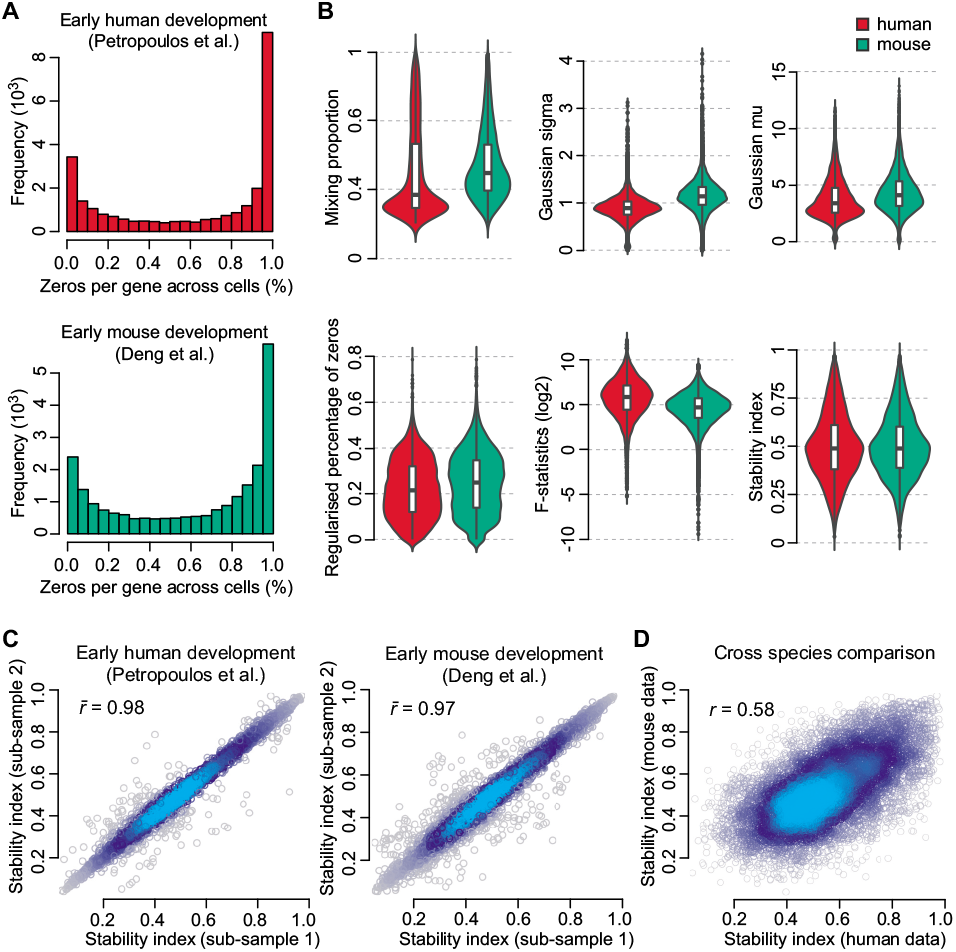
Characterizing gene stability features in single cells from early human and mouse development, respectively. (A) Percentage of zeros per gene across individual cells. (B) Fitted values of mixing proportion (λ), and variance (σ^2^) and mean (μ) in the Gaussian component (top panels) of the mixture model for each gene. Regularised percentage of zeros, *F*-statistics computed from pre-defined cell class and developmental stages (bottom left panel) and stability index derived for each gene for early human and mouse development (bottom right panel), respectively. (C) Scatter plot of stability index calculated from two random sub-sampling of cells from each dataset. Mean Pearson’s correlation coefficient (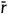) were calculated from pairwise comparison of 10 repeated random sub-sampling on each dataset. (D) Scatter plot of stability index calculated from using the full set of homologous human and mouse genes.

We next investigated the reproducibility of the stability index by randomly sampling 80% of all cells and re-calculating the stability index for each sub-sample. We found the stability index to be highly reproducible in both the human and the mouse data (Figure 2C) with average Pearson correlation coefficients of 0.98 and 0.97. The stability indices also showed relatively high correlation between human and mouse (Figure 2D), suggesting gene expression stability is conserved across species.

### Comparative analysis of SEGs identified in single cells and HKGs defined from bulk transcriptome

To understand the relationships of genes with stable expression in single cells with HKGs defined previously with bulk microarray [16] and RNA-seq [10], we derived a list of SEGs for human and mouse respectively by computing the rank percentiles of stability index as well as the four stability features. Genes with a stability index rank percentile above 80 as well as a reversed rank percentile above 60 for each of the four stability features were included in the SEG list. Using this approach, we derived lists of 1076 human (hSEG) and 830 mouse (mSEG) genes, respectively (Figure 3A and B). In comparison to the HKGs defined previously using bulk transcriptomes, we found that SEGs identified on the single-cell level have significantly smaller expression variances across individual cells (Figure 3A).

**Figure 3.**
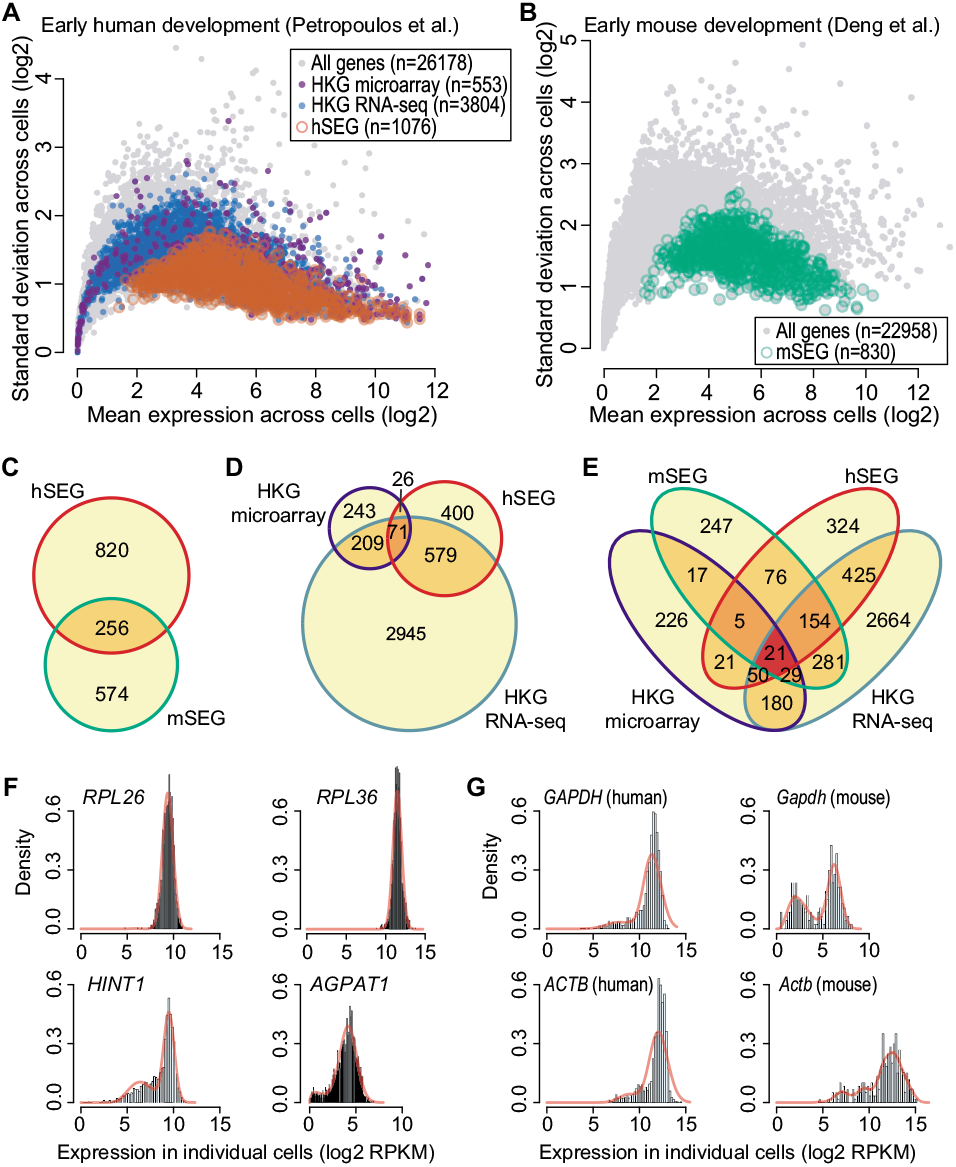
Comparison of SEGs identified on individual cell level using scRNA-seq with HKGs defined on cell population level using bulk transcriptome data. (A) Scatter plot showing mean expression (*x*-axis) and variance (*y*-axis) of each gene (gray circles) across profled single cells. Open red circles represent SEGs identified from early human development data (hSEG) in this study whereas dark and light blue solid circles represent HKGs defined previously using bulk microarray [16] and RNA-seq data [10]. (B) Same as (A) but for SEGs identified from early mouse development data (mSEG; green circles). (C-E) Venn diagrams showing overlaps of SEGs identified from early human and mouse development (C), HKGs defined using bulk microarray and RNA-seq (D), and the overlap of all lists (E). (F) Expression patterns of example genes that are defined as SEGs using scRNA-seq data but not as HKGs using bulk microarray or RNA-seq data (*RPL26* and *RPL36*) and vice versa (*HINT* and *AGPAT*1) across individual cells. (G) Expression patterns for *GAPDH* and *ACTB* in human and mouse (*Gapdh* and *Actb*) across individual cells.

For the human and mouse SEG lists derived from above, there were 256 common genes, accounting for 24% of the hSEG or 31% of the mSEG (Figure 3C). Comparing with previously defined HKGs (Figure 3D), there were 97 common genes between our hSEG list and those defined by microarray (9% and 18%), and 650 between hSEG list and those defined by bulk RNA-seq (60% and 17%). Together, these reflected a relatively low to moderate overlap among SEGs and HKGs (Figure 3E), highlighting both their commonality and the uniqueness, which may be attributed to the biological systems, data resolutions (population vs. individual cells), and analytic approaches from which they are defined.

To investigate the difference between SEGs and HKGs defined by bulk transcriptomes, we inspected a few individual genes that were defined as SEGs using scRNA-seq data but not HKGs by bulk microarray or RNA-seq, and *vice versa.* We discovered that many ribosomal proteins (such as *RPL26* and *RPL36*) that were included in the SEG list but not in the HKG lists (Figure 3F) showed strong unimodal expression patterns across all cells. In contrast, genes such as *HINT1* (Histidine triad nucleotide-binding protein 1) and *AGPAT1* (1-Acylglycerol-3-Phosphate O-Acyltransferase), both of which have been reported to be differentially expressed in brain tissue [41] or malignant oesophageal tissues [42] compared to normal samples, were included in both microarray and RNA-seq defined HKG lists, but not in SEG list due to their bimodal expression patterns across individual cells.

Finally, we examined the expression patterns of *GAPDH* and *ACTB* (Figure 3G), genes which are commonly treated as canonical HKGs for data normalization, and observed clear bimodality in both the human and mouse data. In agreement with previous studies [10,17, 25, 43], these data argue against their usage as “housekeeping genes” for sample normalization.

### SEGs exhibit strong expression stability in single cells across early human and mouse development stages

We hypothesized that if the expression levels of the SEGs are relatively stable, they should show relatively small expression differences across the different cell types from various biological systems. We first investigated principal component analysis (PCA) plots generated from early human and mouse development data using all genes (all expressed mRNA), or subsets of genes defined for human (i.e. HKG microarray, HKG RNA-seq, and hSEG) (Figure 4A) and mouse (i.e. mSEG) (Figure 4B). We found that for human data there is clear separation of developmental stages in the first two principal components when PCA plots were created by using either all genes, and HKGs defined from microarray or RNA-seq, suggesting genes that were expressed differentially in different developmental stages were driving the separation. In contrast, the PCA plot generated from using hSEG show much less separation with respect to the developmental stages, suggesting they are generally expressed at a similar level across individual cells irrespective to cell differentiation and change of developmental stages. Similar results were observed from mouse development data (Figure 4B) where PCA plot generated from mSEG show less cell type and development stage separation compared to PCA plot generated from using all genes.

**Figure 4.**
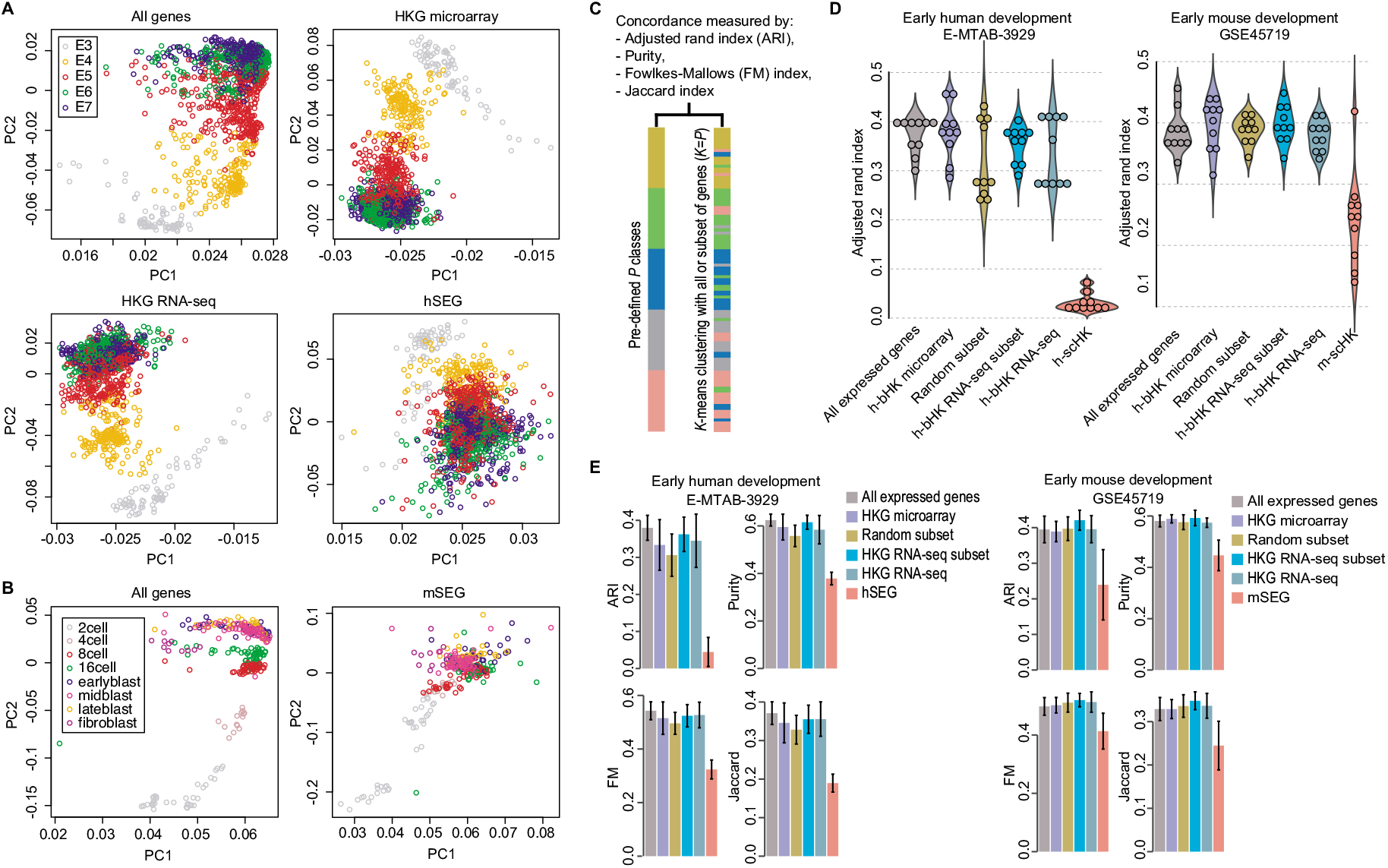
Evaluation of SEGs and HKGs using early human and mouse development scRNA-seq datasets. (A) PCA plots generated from human development data using all expressed genes, HKGs, or hSEGs. Cells are colored by their pre-defined developmental stages. (B) PCA plots generated from mouse development data using all expressed genes or mSEGs. Cells are colored by their pre-defined types and developmental stages. (C) Schematic showing the quantification of concordance of *k*-means clustering with pre-defined cell classes using a panel of metrics. (D) Violin plot of concordance (ARI) between *k*-means clustering and pre-defined cell class labels, using all expressed genes, HKGs, random subsets of genes sampled from all expressed genes, or those from bulk RNA-seq but matched to the size of hSEG and mSEG (HKG RNA-seq subset), respectively. (E) Barplots of concordance between k-means clustering and pre-defined cell class labels, using all expressed genes, genes included in each subset list, and random subsets as in (D) for human and mouse data, respectively.

To quantify the above visual observations in human and mouse developmental datasets, we utilized k-means clustering to partition cells into five and eight clusters respectively, using all genes (all expressed mRNA) or subsets of genes defined in each list (i.e. hSEG, mSEG, HKG microarray and HKG RNA-seq) with the hypothesis that clusters arising from using SEGs and HKGs will exhibit lower concordance with pre-defined cell type- and tissue-specific labels (Figure 4C), thereby demonstrating consistent levels of expression across different cell and tissue types. Random subsets that contained the same number of genes as in SEGs were included by sampling from either all genes or in HKG RNA-seq list to account for the size of the gene-sets used in clustering (see Methods).

Indeed, we found that k-means clustering outputs using SEGs derived from scRNA-seq data showed the lowest concordance to their pre-defined cell class labels (i.e. embryonic day of development or cell types) as quantified by the adjusted rand index (ARI) (Figure 4D) and the three other concordance metrics, namely Purity, Fowlkes-Mallows index (FM), and Jaccard index (Figure 4E). Together, these results demonstrate that SEGs are stably expressed across cells and developmental stages in the two scRNA-seq datasets.

### SEGs maintain expression stability across many different tissues and biological systems

To test whether SEGs derived from the above two early mammalian development datasets are stably expressed in other cell and tissue types, we evaluated these SEGs on eight additional datasets (Table 1) which are independent of the two scRNA-seq datasets used for identifying SEGs. These additional datasets represent drastically different tissues and biological systems in both human and mouse, as well as different sequencing protocols and a wide range in the number of cells sequenced.

Similar to the above section (3.3), we quantified the clustering concordance with respect to each of their pre-defined cell class labels using each of the four concordance metrics (ARI, Purity, FM, and Jaccard) (Table 2). We found that on average, clustering using SEGs gave the lowest concordance to the predefined cell type- and tissue-specific class labels in all tested datasets compared to clustering using all expressed genes or HKGs defined using bulk microarray and RNA-seq datasets. These results suggest that SEGs defined in early human and mouse development also display strong expression stability in various cell/tissue types and biological systems, and they are considerably more stable than HKGs defined using bulk transcriptome data on the single-cell level.

**Table 2.** Stability evaluation results on independent scRNA-seq datasets that profile various cell types and biological systems. All indices are within the range of [0, 1] and are multiplied by 100. The lowest results from each metric in each dataset are bolded.

|  | Peripheral blood mononuclear cells (human); [29] |  |  |  | PSCs and endoderm progenitors (human); [30] |  |  |  |
| --- | --- | --- | --- | --- | --- | --- | --- | --- |
|  | All genes | HKG microarray | HKG RNA-seq | hSEG | All genes | HKG microarray | HKG RNA-seq | hSEG |
| ARI | 55±8 | 42±3 | 38±4 | **29±6** | 69±5 | 58±5 | 55±6 | **41±3** |
| Purity | 69±7 | 62±2 | 59±1 | **52±5** | 80±4 | 74±3 | 71±5 | **59±3** |
| FM | 67±5 | 56±1 | 52±3 | **45±4** | 75±4 | 66±4 | 63±5 | **51±2** |
| Jaccard | 49±6 | 39±1 | 35±2 | **29±4** | 60±5 | 48±4 | 46±6 | **34±2** |
|  | Multicellular metastatic melanoma (human); [31] |  |  |  | Adult and fetal brain (human); [32] |  |  |  |
|  | All genes | HKG microarray | HKG RNA-seq | hSEG | All genes | HKG microarray | HKG RNA-seq | hSEG |
| ARI | 31±5 | 18±2 | 18±1 | **15±1** | 53±7 | 50±3 | 39±4 | **36±3** |
| Purity | 80±5 | 73±1 | 74±1 | **71±1** | 82±3 | 76±4 | 74±3 | **68±2** |
| FM | 51±3 | 39±2 | 40±1 | **37±1** | 62±6 | 59±2 | 50±3 | **47±3** |
| Jaccard | 32±2 | 22±2 | 24±1 | **21±1** | 44±6 | 41±2 | 33±3 | **30±3** |
|  | Cortex and hippocampus (mouse); [33] |  |  |  | Developmental lung epithelial cells (mouse); [34] |  |  |  |
|  | All genes | HKG microarray | HKG RNA-seq | mSEG | All genes | HKG microarray | HKG RNA-seq | mSEG |
| ARI | 45±8 | 36±5 | 31±3 | **27±2** | 61±6 | 55±4 | 48±2 | **45±0** |
| Purity | 72±3 | 66±1 | 63±1 | **58±1** | 83±4 | 80±2 | 76±1 | **74±0** |
| FM | 55±6 | 49±4 | 44±3 | **41±2** | 72±4 | 68±3 | 62±2 | **60±0** |
| Jaccard | 38±6 | 32±4 | 28±2 | **25±1** | 56±5 | 51±3 | 45±2 | **43±0** |
|  | Mesoderm diversification (mouse); [35] |  |  |  | Pancreas inter- and intra-cells (mouse); [36] |  |  |  |
|  | All genes | HKG microarray | HKG RNA-seq | mSEG | All genes | HKG microarray | HKG RNA-seq | mSEG |
| ARI | 54±2 | 43±8 | 49±3 | **32±6** | 37±4 | 22±3 | 23±3 | **18±2** |
| Purity | 66±1 | 62±6 | 65±1 | **59±6** | 89±3 | 78±3 | 76±2 | **72±1** |
| FM | 68±1 | 63±8 | 67±1 | **58±7** | 52±4 | 38±3 | 39±3 | **34±2** |
| Jaccard | 52±1 | 46±7 | 50±1 | **40±8** | 30±3 | 20±3 | 21±3 | **17±2** |

### Gene stability index derived from single cells correlates with gene sequence and structural characteristics

To further characterize gene expression stability in single cells, we correlated the stability index and each stability feature extracted from scRNA-seq data with various gene structural and conservation features calculated from various data sources. We found that the stability index correlated positively with the number of exons in a gene, gene expression, and gene conservation, and negatively with GC-content in the gene body in both human and mouse (Figure 5A), many of which are characteristics of HKGs reported in previous studies. Consistent with this, we found SEGs are more evolutionarily conserved [44] with higher phyloP scores. SEGs also possess more exons, in agreement with previous finding on HKGs [45], despite mouse genes on average having fewer exons than human genes. Both human and mouse SEGs appeared to have a slightly lower GC-content but, similar to previous observation on HKGs, the relation was relatively weak [46] (Figure 5B).

**Figure 5.**
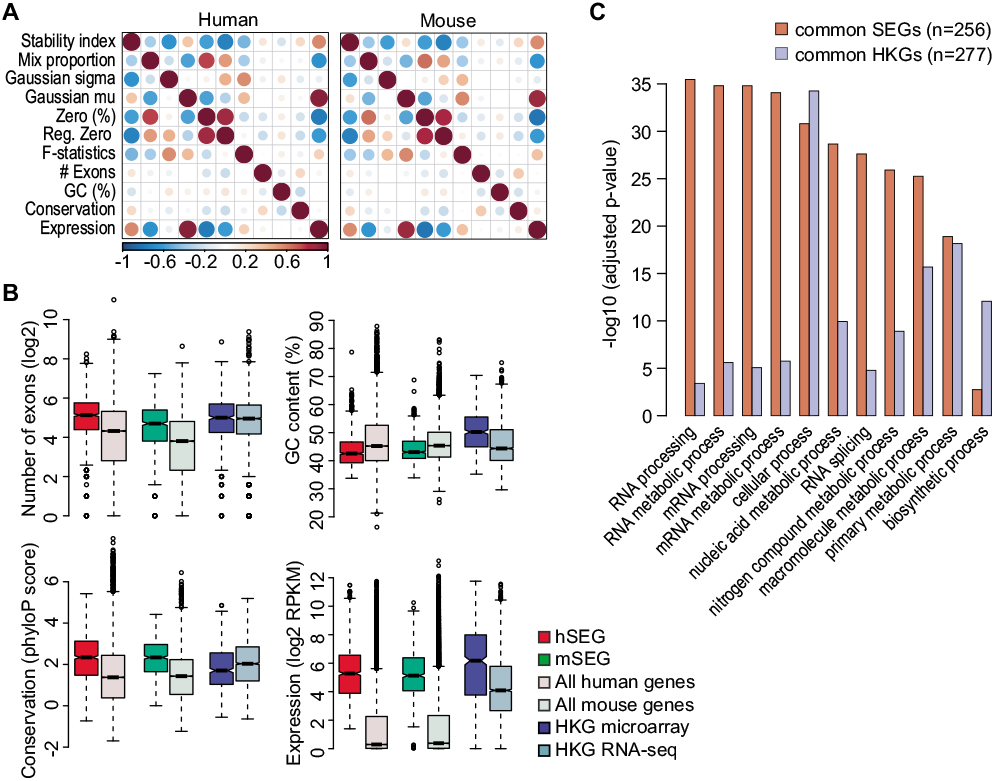
Characterization of stability index with sequence and gene characteristics. (A) Pearson correlation analyses of human and mouse gene stability features with respect to genomic structural and evolutional gene features. (B) Boxplots of various gene characteristics for SEGs, HKGs and all expressed genes. (C) Gene ontology analysis (by over-representation) of SEGs that are common between hESG and mSEG; and HKGs that are common between HKG microarray and HKG RNA-seq.

Perhaps unsurprisingly, SEGs identified in this study possess similar characteristics to those observed in HKGs, indicating that they are serving essential cellular functions akin to HKGs. Supporting this, we found that multiple top-enriched Gene Ontology terms that describe essential cellular functions are shared by common SEGs (genes overlap between hESG and mSEG) and common HKGs (genes overlap between HKG microarray and HKG RNA-seq) (see Methods for details). Nevertheless, common SEGs are far more enriched for most Ontology terms than common HKGs defined from bulk transcriptome, indicating the higher resolution enabled by scRNA-seq data for identifying genes that are truly stably expressed across individual cells.

## Discussion

Since the emergence of high-throughput transcriptome profiling, the search for stably expressed genes (SEGs) has been a central quest in modern biology. Such genes are often thought to be essential for basic cellular functions given their relatively constant expression and activity despite changes in cell status and types. The hypothesis that such genes may serve the same housekeeping functions across various cell and tissue types has also led to their definition as “housekeeping genes” (HKGs). While the existence of true HKGs whose expression are universally constant across all cells and systems is a subject of debate [42, 47], their practical usage as control genes for experimental data normalization is well appreciated.

Recent advances in single-cell transcriptome profiling using scRNA-seq have highlighted the phenomenal amount of gene expression stochasticity and heterogeneity in single cells. Compared to bulk transcriptome data that aggregate millions of cells to obtain a single gene expression measure, scRNA-seq data allows the expression dynamics of each gene within individual cells to be monitored, and therefore enables the identification of genes that are truly expressed at a steady level in individual cells across tissues and developmental stages. By modeling from large-scale scRNA-seq datasets, we quantified the relative expression stability of genes on the single-cell levels. We showed that the SEGs derived based on their stability indices are considerably more stable in not only the scRNA-seq datasets from which they are identified but also independent scRNA-seq datasets that profiles various cell types and biological systems.

Our analysis demonstrated that despite the high variability in single-cell gene expression, a subset of genes is inherently more stable in expression than other genes within individual cells. Their sequence and gene structural properties are strongly reminiscent of HKGs defined from bulk transcriptome, suggesting their essential roles in maintaining basic cellular functions on the individual cell level.

The proposed framework can be applied in a data dependent manner to rank genes based on their expression stability in a given scRNA-seq dataset. This relaxes the rigid binary definition of HKGs and enable a more practical definition of stable expression in different experimental contexts. Hence, the proposed method is particularly useful for defining stable or “control” genes in various scRNA-seq experiment, which is often a key step in normalizing such data [48, 49].

The generalizability of SEGs is dependent on the diversity of cell types profiled in a scRNA-seq experiment. Various cell atlas profling initiatives such as the Human Cell Atlas (https://www.humancellatlas.org) is currently under way to comprehensively characterize the transcriptome of every human cell. Information from such resources in conjunction with our computational framework will provide an even more precise assessment of gene expression stability in single cells that will enrich subsequent avenues of research including characterizing heterogeneity and stability of single-cell transcriptomes and their use for technical data normalization and standardization.

Taken together, this comprehensive evaluation study demonstrates the utility of measuring gene expression stability at the single-cell level and marks a shift in paradigm for selecting genes that are stably expressed in single cells for practical applications.

## Methods

### Evaluating the stability of gene lists

To assess the expression stability of each gene list in various cell types and biological systems, the k-means algorithm was utilized to cluster each scRNA-seq data to its pre-defined number of clusters and an array of evaluation metrics were applied to compute the concordance with respect to the pre-defined (“gold standard”) class labels. Evaluation metrics include the adjusted Rand index (ARI), Purity, the Fowlkes-Mallows index (FM) and the Jaccard index.

Let *U* = {*u*_1_, *u*_2_,…,*u_P_*} denote the true partition across *P* classes and *V* = {*v*_1_,*v*_2_,…,*v_k_*} denote the partition produced from k-means clustering (*K = P*). Let *a* be the number of pairs of cells correctly partitioned into the same class by the clustering method; *b* be the number of pairs of cells partitioned into the same cluster but in fact belong to different classes; *c* be the number of pairs of cells partitioned into different clusters but belongs to the same class; and *d* be the number of pairs of cells correctly partitioned into different clusters. Then the Adjusted Rand Index [50], the Jaccard index [51], and the Fowlkes-Mallows index [52] can be defined as and the Purity [53] can be calculated as

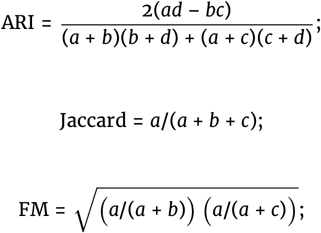

and the Purity [53] can be calculated as

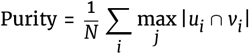

where *N* is the total number of cells, *i* and *j* are the indices of clusters from clustering output *u_i_* and pre-defined class label *v_j_*.

For each dataset, we calculated and compared the above four metrics using (i) all expressed genes, (ii) HKGs defined using microarray data [16], (iii) HKGs defined using bulk RNA-seq data [10], and (iv) SEGs identified in this study. In order to account for potential effects of gene list length, we also generated random subsets with the same number of genes in our SEG lists first by randomly sampling from all expressed genes in the dataset, and second by randomly sampling from the HKG list defined by bulk RNA-seq. Since the k-means clustering algorithm is not deterministic and the random sampling process introduces variability, the above procedure was repeated 10 times to account for such variability.

### Gene properties

To characterize SEGs identified in early human and mouse development datasets, we extracted gene sequence and structural features including the number of exons and percentage GC content in the gene body for human and mouse, respectively, using the biomaRt [54]. Additionally, to characterize gene evolutionary conservation, phyloP scores were downloaded from the UCSC Genome Browser for mouse (mm10) and human (hg38) genomes. Exonic bases of each gene were determined based on GENCODE Genes for human (release 26) and mouse (release 14). The set of conservation scores for each gene was averaged for each gene. We assessed the concordance of gene expression stability index and each stability feature derived from single cells with structural features, conservation scores, and their expression across all genes for human and mouse using Pearson correlation coefficients. We also compared these features for SEGs and previously defined HKGs against all expressed genes in human and mouse, respectively.

### Gene ontology enrichment analysis

To perform gene ontology enrichment analysis, we first defined SEGs that are shared between hSEG and mSEG as “common SEGs” and HKGs that are shared between HKG microarray and HKG RNA-seq as “common HKGs”. The similar numbers of common SEGs (256) and common HKGs (277) allowed us to avoid the gene-set size bias in the enrichment analysis.

Over-representation of common SEGs or common HKGs was evaluated by comparing each set of genes against ontologies defined in Gene Ontology database [55]. Fisher’s exact test was used to assess statistical significance. Top-enriched ontologies from either common SEGs or common HKGs were combined for interpretation.

## Availability of supporting data and materials

The datasets generated and/or analyzed during the current study are available in either the NCBI GEO repository or the EMBL-EBI ArrayExpress repository (Table 1). An interactive web resource is available at http://shiny.maths.usyd.edu.au/SEGs. The computational framework implemented in R is available from https://github.com/PengyiYang/SEGs.

## Declarations

## List of abbreviations

scRNA-seq: : Single-cell RNA-seq
HKGs: : housekeeping genes
SEGs: : stably expressed genes
SAGE: : serial analysis of gene expression
hSEG: : stably expressed genes derived from early human developmental dataset
mSEG: : stably expressed genes derived from early human developmental dataset
HKG microarray: : housekeeping genes defined using bulk microarray
HKG RNA-seq: : housekeeping genes defined using bulk RNA-seq
PCA: : principal component analysis
ARI: : adjusted rand index
FM: : Fowlkes-Mallows index.

## Consent for publication

Not applicable

## Author’s Contributions

PY conceived the study with input from JYHY. All authors contributed to the design, analytics, interpretation and the direction of the study. YL and PY lead the analytics and AYW lead the curation of the datasets. All authors wrote, reviewed, edited, and approved the final version of the manuscript.

## Acknowledgements

The authors thank their colleagues at the School of Mathematics and Statistics, The University of Sydney, and Prof. Ze-Guang Han and Dr. Xianbin Su at Shanghai Jiao Tong University for informative discussion and valuable feedback.

